# ST8Sia2 polysialyltransferase protects against infection by *Trypanosoma cruzi*

**DOI:** 10.1101/2023.11.14.567071

**Authors:** Bruno Rafael Barboza, Janaina Macedo-da-Silva, Lays Adrianne Mendonça Trajano Silva, Vinícius de Morais Gomes, Deivid Martins Santos, Antônio Moreira Marques Neto, Simon Ngao Mule, Juliana Borsoi, Carolina Borsoi Moraes, Martina Mühlenhoff, Walter Colli, Suely K. N. Marie, Lygia da Veiga Pereira, Maria Julia Manso Alves, Giuseppe Palmisano

**Author notes:** Correspondence to: Giuseppe Palmisano, Department of Parasitology, Institute of Biomedical Sciences, University of São Paulo, Avenida Lineu Prestes 1374, São Paulo, Brazil.

## Abstract

Glycosylation is one of the most structurally and functionally diverse co- and post-translational modifications in a cell. Addition and removal of glycans, especially to proteins and lipids, characterize this process which have important implications in several biological processes. In mammals, the repeated enzymatic addition of a sialic acid unit to underlying sialic acids (Sia) by polysialyltransferases, including ST8Sia2, leads to the formation of a sugar polymer called polysialic acid (polySia). The functional relevance of polySia has been extensively demonstrated in the nervous system. However, the role of polysialylation in infection is still poorly explored. Previous reports have shown that *Trypanosoma cruzi* (*T. cruzi*), a flagellated parasite that causes Chagas disease (CD), changes host sialylation of glycoproteins. To understand the role of host polySia during *T. cruzi* infection, we used a combination of *in silico* and experimental tools. We observed that *T. cruzi* reduces both the expression of the ST8Sia2 and the polysialylation of target substrates. We also found that chemical and genetic inhibition of host ST8Sia2 increased the parasite load in mammalian cells. These findings suggest a novel approach to interfere with parasite infections through modulation of host polysialylation.

**AUTHOR SUMMARY:** Glycosylation is a co- and/or post-translational modification regulated by the addition and removal of glycans. This process shapes the cellular glycome, which in turn, holds significant implications in various biological processes. *Trypanosoma cruzi* (*T. cruzi*), the etiological agent of Chagas disease, a globally concerning neglected tropical disease affecting 6 to 8 million individuals worldwide, exerts a profound influence on host glycoprotein sialylation. Remarkably, *T. cruzi* is incapable of synthesizing sialic acid (Sia) and relies on acquiring it from host glycoconjugates. In mammals, the formation of polysialic acid (polySia) is mediated by polysialyltransferases, such as ST8Sia2. The functional relevance of polySia has been extensively documented in the nervous system. Nevertheless, its role within the context of infectious processes remains largely unexplored. Herein, we demonstrate that in *T. cruzi*-infected host cells, the expression of the ST8Sia2 enzyme is downregulated, resulting in diminished levels of polysialylation. Furthermore, a reduction in the levels of NCAM1 and SCN5A was observed, which can be attributed to the decreased host polysialylation. Moreover, enzymatic removal of polySia, along with chemical inhibition and genetic silencing of ST8Sia2, led to a marked increase in the number of intracellular parasites. We posit that ST8Sia2 inhibition favors *T. cruzi* infection, thereby elucidating novel avenues for understanding the mechanisms associated with Chagas disease pathogenesis, prominently featuring the pivotal role of host polysialylation.

## INTRODUCTION

Glycans are important regulators in different biological processes, with prominent roles in cell adhesion [1], cell activation and signaling [2,3], modulation of immune responses [4,5], cell death [6], and initiation of tumor progression and metastases [7]. The addition and removal of glycans (mono, oligo and polysaccharides) to proteins, lipids [8] and small RNAs [9] is performed by an array of glycosyltransferases and glycosidases, that act in a concerted way to remodel the cellular glycome [10,11]. Glycosylation is one of the most structurally and functionally complex co- and/or post-translational modifications that occur in cells [12,13]. Glycans attached to glycoconjugates, commonly expressed on the surface of cells, exhibit enormous variability and structural complexity, contributing to their respective biological activities [14].

The majority of glycan chains are composed of monosaccharides with five- or six-carbons, with the striking exception of sialic acid (Sia), which belongs to a large family of sugars with a nine-carbon atom backbone and typically found attached to the terminal position of glycans [15]. The glycosidic linkage between a Sia monomer to another underlying Sia residue generates homo-oligo/polymer structures from di- to polysialic acid (polySia) with different α2,4/8/9 intersialyl linkages [16,17]. The biosynthesis of α2,8-linked polySia is catalyzed by ST8Sia polysialyltransferases (polySTs) [18]. In mammals, polysialylation is catalyzed by Golgi-resident polySTs: ST8Sia2 and ST8Sia4 [16,19]. Recently, ST8Sia3 was reported as a polyST capable of autopolysialylation [20]. Structurally, polySia is characterized by repeated Sia monomers with a degree of polymerization ranging from 8 to 400 sialic acid molecules [21]. The presence of polySia is confined to a restricted set of glycoproteins, with the neural cell adhesion molecule 1 (NCAM1) as the best-studied polySia-carrier in mammals [22].

The biological relevance of polySia in mammals has been extensively explored in the brain, highlighting its role in the migration of neuronal precursor cells [23], synaptic neuronal plasticity [19], and axonal outgrowth [24]. The involvement of polySia has also been demonstrated in the development of other organs, including the liver [25], kidney [26], placenta [27], and heart [28]. Beyond the considerable number of studies that have provided important information about the role of polysialylation in the nervous system, the participation of polySTs and polySia in the cardiac system has been less explored. Recently, NCAM and polysialylation have been shown to play critical roles in the cardiac conduction system [29]. Additionally, in ST8Sia2 knockout mice, polysialylation deficiency compromised the action potential and the voltage-gated sodium channel functions in atrial cardiomyocytes [30], experimentally evidencing that modulation in polysialylation is sufficient to alter the complex developmental biology and function of cardiomyocytes.

Chagas disease, also known as American trypanosomiasis, is an anthropozoonosis caused by the protozoan *T. cruzi* that affects between 6 and 8 million individuals worldwide, being considered a neglected tropical disease in the world [31,32]. Clinically, CD can be characterized by two main successive phases, the acute and the chronic phases [33]. Among the clinical manifestations of Chagas disease, Chronic Chagas Cardiomyopathy (CCC), presents itself as the most severe manifestation of the disease and affects 1/3 of infected individuals. CCC comprises four main syndromes: *heart failure, arrhythmias, thromboembolism* and *anginal manifestations* that can simultaneously occur in the same individual [34,35].

In the course of host infection, *T. cruzi* uses a wide variety of mechanisms to escape from host immune surveillance and establish a successful infection, with emphasis on *Trans*-sialidases (TS) [36,37] which catalyze direct Sia transfer between macromolecules, a novel enzymatic activity that was earlier demonstrated [38]. During host cell adhesion and invasion, these enzymes provide an elegant mechanism in which *T. cruzi* specifically captures α2,3-sialic acid from host cells and covers its own surface molecules, mainly mucins, as reviewed in [37,39,40]. Of note, *T. cruzi* parasites are incapable of synthesizing Sia [41,42]. In fact, parasite cells transfer Sias from host cell-surface, the internal surface of the phagolysosome of host cells and from serum sialoglycoproteins to trypanosome cell-surface glycoconjugates [36,43,44].

Previous studies reported that in *T. cruzi* infection, possible disorders of host aberrant glycosylation are due to impairment of glycoprotein sialylation. In 1986, Libby and colleagues [45] demonstrated *in vitro* that *T. cruzi* trypomastigotes modify the surface of rat myocardial and human vascular endothelial cells by desialylation. According to Marin-Neto and colleagues [46], the loss of Sias on the surface of endothelial cells can promote an increase in platelet aggregation and, consequently, microvascular thrombosis. More recently, it was demonstrated that CD43, a sialoglycoprotein expressed at high levels on leucocytes and previously reported as a natural receptor for *T. cruzi* trans-sialidase [47], plays a critical role in the pathogenesis of acute chagasic cardiomyopathy. In this study, a reduced cardiac parasite load in CD43 knockout mice [48] was observed, suggesting the involvement of CD43 in the development of Chagas cardiomyopathy.

Considering the clinical complications related to Chagas cardiomyopathy, and the limited knowledge on the involvement of cardiac glycosylation in susceptibility and/or resistance to *T. cruzi* infection, the interplay between the modulation of glycosylation and possible changes in the pathophysiology of Chagas cardiomyopathy represents an interesting target to explore. In the present study, we report new molecular mechanisms used by *T. cruzi* to establish a successful infection on host cells. The current study evaluated the modulation of host polysialylation on experimental *T. cruzi* infection using a combination of *in silico* and experimental tools. We demonstrated that *T. cruzi* infection significantly reduces ST8Sia2 levels and substrate polysialylation. We also verified that the chemical and genetic inhibition of ST8Sia2 increases parasite load in host cells, suggesting a functional role of host polysialylation during infection by *T. cruzi*.

## RESULTS

### Modulation of genes involved in host N-glycosylation machinery by *T. cruzi* is accompanied by downregulation of ST8Sia2 polysialyltransferase

In this study, we first performed an *in-silico* analysis on publicly available transcriptome data obtained from human induced pluripotent stem cell-derived cardiomyocytes (hiPSC-CM) infected with *T. cruzi* Y strain [49] to map the differential expression of glycan-related genes (glycogenes) **(Supplementary Figure S1)**. A total of 39 glycogenes, including α-glucosidases, fucosyltransferases, galactosyltransferases, glucosyltransferases, mannosidases, mannosyltransferases, n-acetylglucosaminyltransferases, sialidases, and sialyltransferases were modulated in infected hiPSC-CM **(Supplementary Figure S1A)**. Among the regulated glycogenes, ST8 alpha-N-acetyl-neuraminide alpha-2,8-sialyltransferase 2 (ST8Sia2) was downregulated in *T. cruzi-*infected hiPSC-CM **(Supplementary Figure S1B–D)**, being more pronounced at 48 hours post-infection (h.p.i.). A previous study also showed that polysialylation deficiency compromises the functionality of atrial cardiomyocytes in ST8Sia2 knockout mice [30]. Taken together, these findings strongly suggested that in hiPSC-CM, *T. cruzi* parasite modulates the expression of glycogenes, and manipulates host polysialylation machinery by downregulating ST8Sia2.

### *T. cruzi* infection modulates hiPSC-CM ST8Sia2 expression and compromises host polysialylation

In order to investigate the impact of host polysialylation upon *T. cruzi* infection, we differentiated human pluripotent stem cells (hiPSC) into cardiomyocytes (hiPSC-CM), and the efficiency of differentiation on hiPSC-CM was determined by the expression of troponin T and α-actinin, using immunofluorescence microscopy **(Supplementary Figure S2A),** and by the percentage of positive cells stained with anti-troponin T (TNNT2+)antibodies analyzed by flow cytometry **(Supplementary Figure S2B)**. Then, we evaluated the levels of polysialylation in hiPSC-CM infected with *T. cruzi* (**Figure 1**). *T. cruzi* parasites were efficiently detected 48 h.p.i. (**Figure 1A**), as described previously [50]. Corroborating our initial data, we observed a 70% downregulation of ST8Sia2 mRNA in *T. cruzi-*infected hiPSC-CM using qRT-PCR (**Figure 1B**). Furthermore, ST8Sia2 protein levels are also reduced in infected hiPSC-CM (**Figure 1C, G)**.

**Figure 1.**
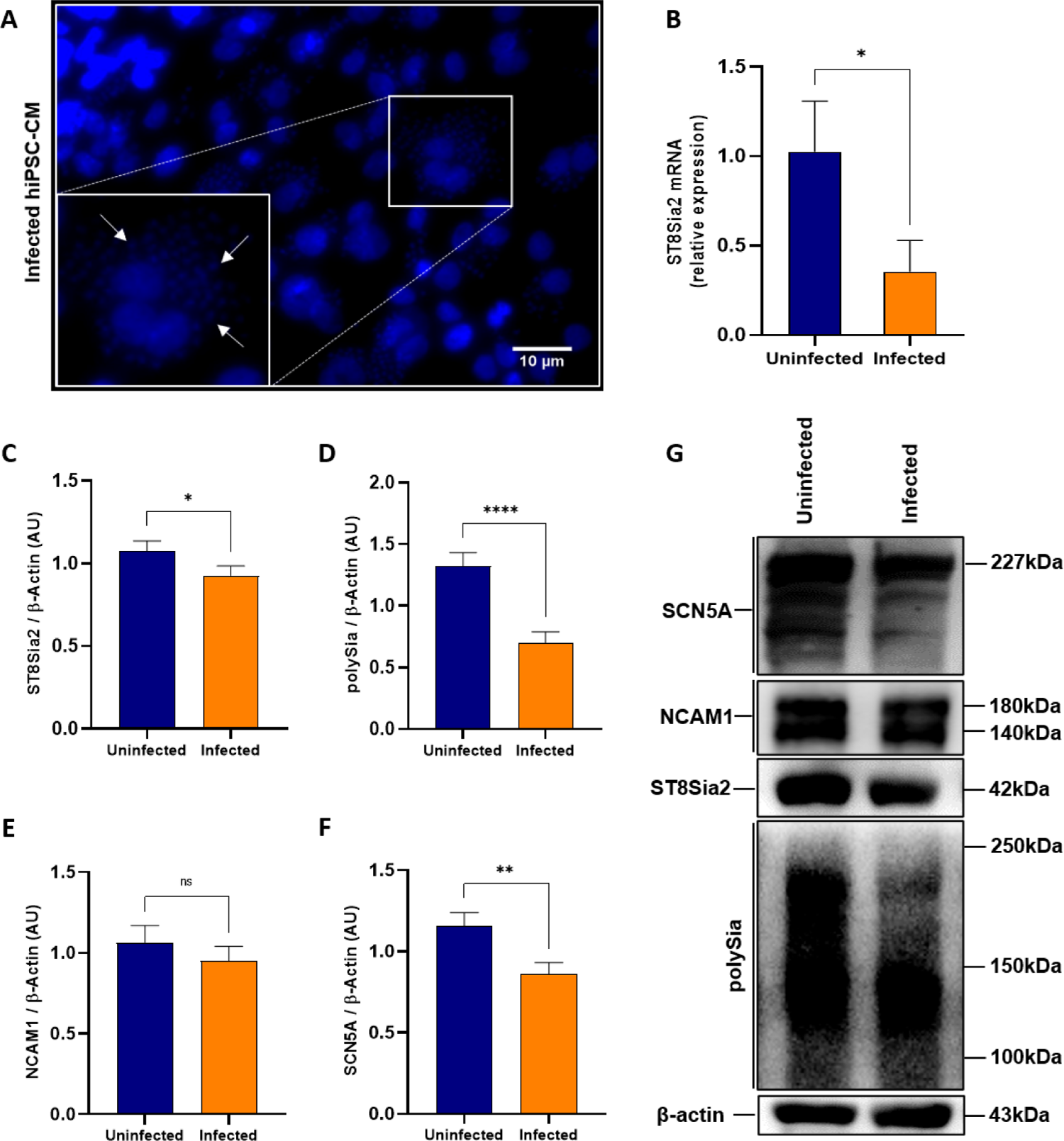
*T. cruzi* infection modulates the abundance of ST8Sia2, SCN5A, and polysialylation on hiPSC-CM. **(A-G)** hiPSC-CM cells were seeded in 24-well microplates (2 x 10^5^ cells/well) and infected with *T. cruzi* trypomastigotes (Y strain) at ratio of 1:5 (hiPSC-CM:trypomastigotes), follow by 48 h incubation at 37 °C. hiPSC-CM cells incubated with medium alone (Uninfected) were used as negative control; **(A)** Representative immunofluorescence images of *T. cruzi-*infected hiPSC-CM. Cell nucleus were stained with DAPI (scale bars = 10 µm). White arrows are representative of *T. cruzi* amastigotes inside hiPSC-CM cells; **(B)** Relative expression of mRNA *ST8Sia2* measured by qRT-PCR in *T. cruzi-*infected hiPSC-CM. The Ct values of the target transcripts were normalized to the relative expression of *GAPDH* as endogenous control, and the relative expression of *ST8Sia2* transcripts was quantified by the 2^-ΔΔ^ Ct method. Each bar represents the mean of three independent experiments performed in triplicate; **(C-G)** Representative Western blotting quantification of ST8Sia2 **(C)**, polySia **(D)**, NCAM1 **(E)**, and SCN5A **(F)** in *T. cruzi-*infected hiPSC-CM cells; **(G)** Representative Western blotting images of levels of ST8Sia2 **(C)**, polySia **(D)**, NCAM1 **(E)**, and SCN5A **(F)** in *T. cruzi-*infected hiPSC-CM cells. The protein levels were analyzed by Western blotting of RIPA cell lysates (15 µg protein) performed under reducing conditions. Results are presented as arbitrary densitometry units (AU). After normalization to the corresponding β-actin content (endogenous control), data were plotted as the ratio between the values obtained in infected and uninfected cells in their respective endogenous controls. Each bar represents the mean of three independent experiments performed in triplicate. Results are expressed as mean ± SEM. Significant differences compared to the uninfected cells are shown by (*) *p* < 0.05, (**) *p* < 0.001, and (***) *p* <0.0001; ns = not significant.

Since polysialylation is a post-translational modification restricted to a set of glycoproteins [16], including the Sodium channel protein type 5 subunit alpha (SCN5A) of the primary cardiac isoform of the voltage-gated sodium channel (Na_V_1.5), and neural cell adhesion molecule 1 (NCAM1) [51], both involved in modulating cardiac functions [29,52], both molecules were focused herein. Following the reduction in ST8Sia2 levels observed in Figure 1, reduced polySia levels were also detected in *T. cruzi-*infected hiPSC-CM (Figure 1D,G). No significant modulation in NCAM1 levels was observed between hiPSC-CM infected and non-infected with *T. cruzi* (Figure 1E,G), as expected since the cardiomyocyte model used in this study did not express PSA-NCAM, the polysialylated form of NCAM1 **(Supplementary Figure S2C)**. Accordingly, no signal was detected using a specific antibody for PSA-NCAM in non-infected **(Supplementary Figure S2C; lane 1)** and infected **(Supplementary Figure S2C; lane 2)** hiPSC-CM. As controls, the efficiency of immunostaining was demonstrated in TE671 cells **(Supplementary Figure S2C; lane 3)**, a human rhabdomyosarcoma cell previously described as positive for NCAM and PSA-NCAM [53] and the specificity of the antibody towards host polysialylation was demonstrated using *T. cruzi* trypomastigotes **(Supplementary Figure S2C; lane 4)**. In addition, we detected a reduction in SCN5A levels in *T. cruzi-*infected hiPSC-CM (Figure 1F**, G)**. These data strongly suggested that infection of hiPSC-CM with *T. cruzi* leads to the downregulation of ST8Sia2, affecting polySia transfer to specific substrates such as SCN5A, thereby reducing their abundance.

### *T. cruzi* infection reduces the expression of host polysialylated target molecules

In order to broaden the approach to the effect of *T. cruzi* infection on the host polysialylation machinery, it became interesting to investigate the impact on polysialylation in SH-SY5Y cells, a human cell line of neuroblastoma. The infection efficiency of SH-SY5Y cells with *T. cruzi* was evaluated after 48 h.p.i by quantification of internalized parasites (Figure 2A). Next, we investigated the impact of *T. cruzi* infection on the polysialylation of SH-SY5Y cells (Figure 2B-G). Similar to the findings observed on *T. cruzi-*infected hiPSC-CM, we detected that ST8Sia2 (Figure 2B**, C)**, polySia (Figure 2B**, D)**, and SCN5A (Figure 2B**, F)** levels were significantly reduced in *T. cruzi-*infected SH-SY5Y cells. We also observed reduced levels of NCAM1 (Figure 2B**, E)** in SH-SY5Y cells infected with *T. cruzi*. Furthermore, there was a significant reduction in the abundance of PSA-NCAM (Figure 2B**, G)**. Thus, these findings reinforce our hypothesis that the polysialylation of host cells is modulated during the course of *T. cruzi* infection.

**Figure 2.**
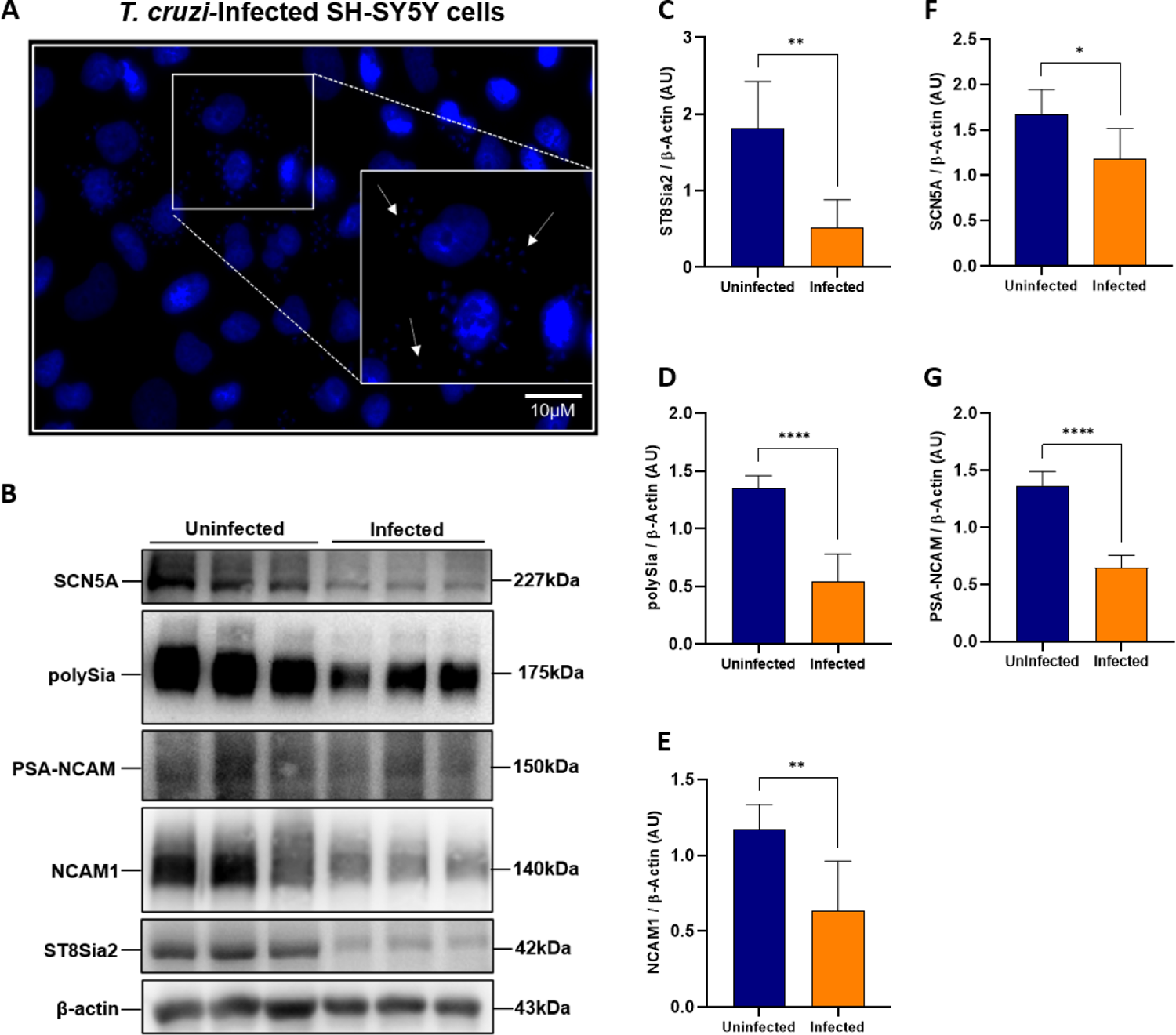
*T. cruzi* infection modulates the polysialylation on SH-SY5Y cells and affects the abundance of polysialylated molecules. (A) Representative immunofluorescence images of *T. cruzi-* infected SH-SY5Y cells. SH-SY5Y cells were seeded in 24-well microplates (5 x 10^5^ cells/well) and infected with *T. cruzi* trypomastigotes (Y strain) at ratio of 1:5 (SH-SY5Y:trypomastigotes) at 37 °C. 48 h.p.i SH-SY5Y cells incubated with medium alone (Uninfected) was used as negative control. Cell nucleus were stained with DAPI (scale bars = 10 µm). White arrows are representative of *T. cruzi* amastigotes inside in SH-SY5Y cells; **(B)** Representative Western blot images of ST8Sia2 levels **(C),** polysialic acid (polySia) **(D)**, NCAM1 **(E)**, SCN5A (**F)**, and PSA-NCAM **(G)** in *T. cruzi-*infected SH-SY5Y cells. SH-SY5Y cells were seeded in 6-well microplates (1 x 10^6^ cells/well) and infected with *T. cruzi* as in **A**. 48 h.p.i, the proteins levels were analyzed by Western blotting of RIPA cell lysates (15 µg protein) performed under reducing conditions; **(C-G)** Western blotting quantification of ST8Sia2 **(C)**, polySia **(D)**, NCAM1 **(E)**, SCN5A **(F)**, and PSA-NCAM **(G)** presented as arbitrary densitometry units (AU). After normalization to the corresponding β-actin content (endogenous control), data were plotted as the ratio between the values obtained in infected and uninfected cells in their respective endogenous controls. Each bar represents the mean of three independent experiments performed in triplicate. Results are expressed as mean ± SEM. Significant differences compared to the uninfected cells are shown by (*) *p* < 0.05, (**) *p* < 0.001, and (****) *p* < 0.0001.

Furthermore, we evaluated the abundance of ST8Sia2 and polySia in *T. cruzi*-infected SH-SY5Y cells using immunofluorescence microscopy approach. Corroborating the data presented in Figure 2, we observed a significant reduction in the abundance of ST8Sia2 (Figure 3A**, C)** and polySia (Figure 3B**, D)** in *T. cruzi-*infected SH-SY5Y cells, when compared to non-infected cells. Next, we quantified the polySia content in the supernatant of SH-SY5Y cells infected with *T. cruzi* using a chromatographic method with fluorescence-based detection. For this assay, SH-SY5Y cells infected and non-infected with *T. cruzi* were incubated with endo-neuraminidase (EndoN), an enzyme that specifically cleaves α2,8-linked sialic acid polymers [16]. We observed that the polySia levels from the supernatant of *T. cruzi-*infected SH-SY5Y cells were significantly reduced compared to non-infected SH-SY5Y cells (Figure 3E), reinforcing our findings that host polysialylation is differentially modulated upon *T. cruzi* infection. In addition, we treated SH-SY5Y cells with EndoN to remove polySia and evaluated whether it would have an impact on *T. cruzi* infection, adopting the workflow shown in Figure 4A. Our results showed that removal of polySia in SH-SY5Y cells increased the number of intracellular parasites 48 h.p.i (Figure 4B,C), reinforcing that host polysialylation has a vital role in the course of *T. cruzi* infection.

**Figure 3.**
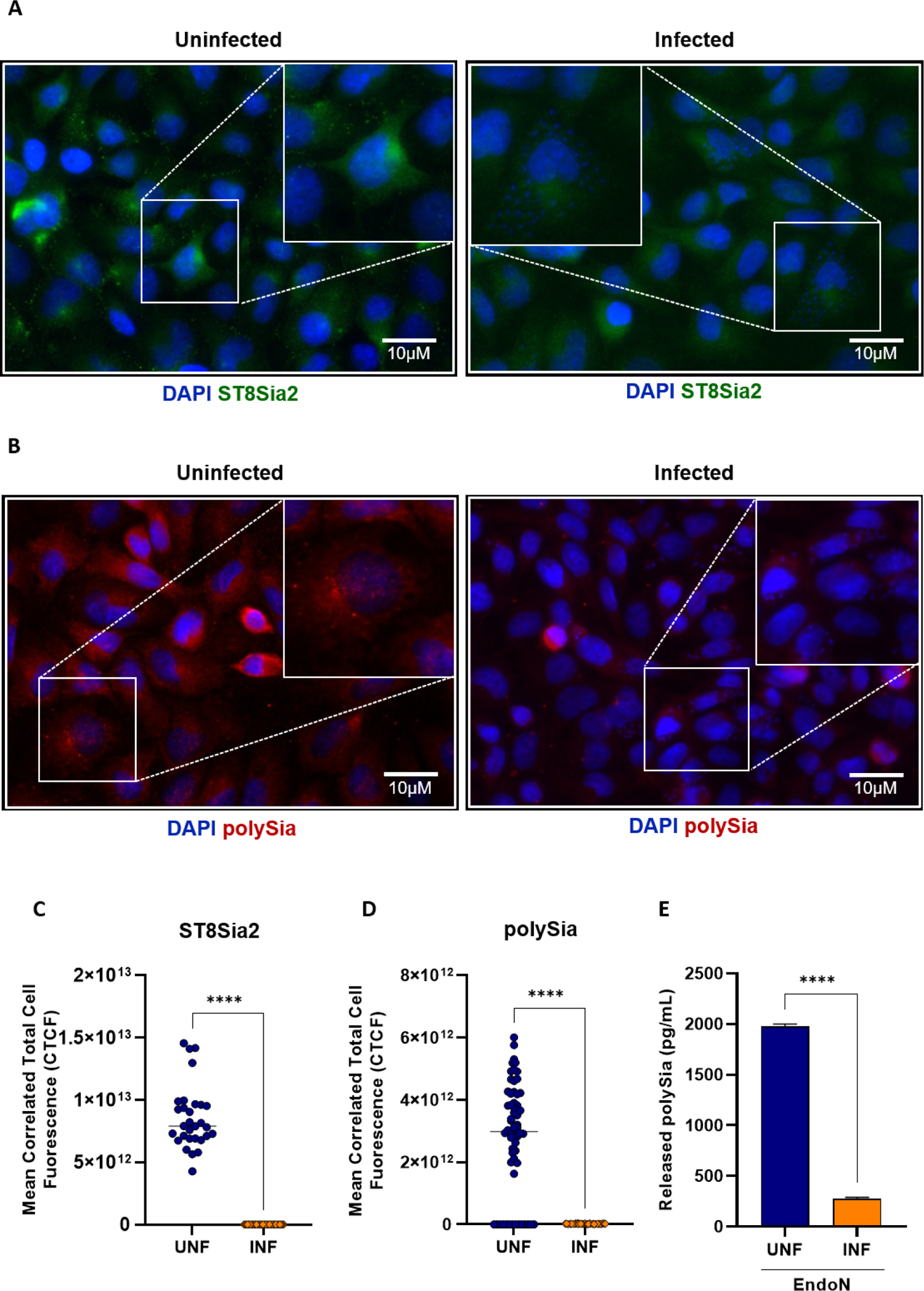
*T. cruzi* infection reduces ST8Sia2 and PolySia levels in SH-SY5Y cells. **(A,B)** Representative images of ST8Sia2 **(A)** and PolySia **(B)** levels in *T. cruzi-*infected SH-SY5Y cells. SH-SY5Y cells were seeded in 24-well microplates (5 x 10^5^ cells/well) and infected with *T. cruzi* trypomastigotes (Y strain) at ratio of 1:5 (SH-SY5Y:trypomastigotes) at 37 °C. After 48 h.p.i, *T. cruzi-* infected SH-SY5Y cells were stained with specific anti-ST8Sia2 **(A)** and anti-polySia **(B)** antibodies, and labeling for the targets was visualized by immunofluorescence microscopy. SH-SY5Y cells incubated with medium alone (Medium) was used as negative controls. Cell nucleus were stained with DAPI (scale bars = 10 µm); **(C,D)** Quantification of ST8Sia2 **(C)** and polySia **(D)** fluorescence intensity in *T. cruzi-*infected SH-SY5Y cells using the calculation for corrected total cell fluorescence (CTCF) as explained in Material and Methods section. Each dot represents the CTCF read out from one cell. A total of 50 SH-SY5Y cells per condition (infected and uninfected) were quantified. The values are expressed as Mean CTCF ± standard error of the mean (SEM); **(E)** polySia levels measured in *T. cruzi*-infected and non-infected SH-SY5Y cells supernatants. SH-SY5Y cells were seeded in T75-flasks (5 x 10^5^ cells/mL) and infected with *T. cruzi* trypomastigotes (Y strain) at ratio of 1:5 (SH-SY5Y:trypomastigotes) at 37 °C. After 48 h.p.i, *T. cruzi-* infected SH-SY5Y cells were treated with EndoN [0.5 µg/ml] for 1h at 37 °C, and polySia levels were measured in supernatants using UHPLC. The same procedure was applied to non-infected SH-SY5Y cells. The results are expressed in pg/mL. Significant differences compared to the uninfected are shown by (****) *p* < 0.0001.

**Figure 4.**
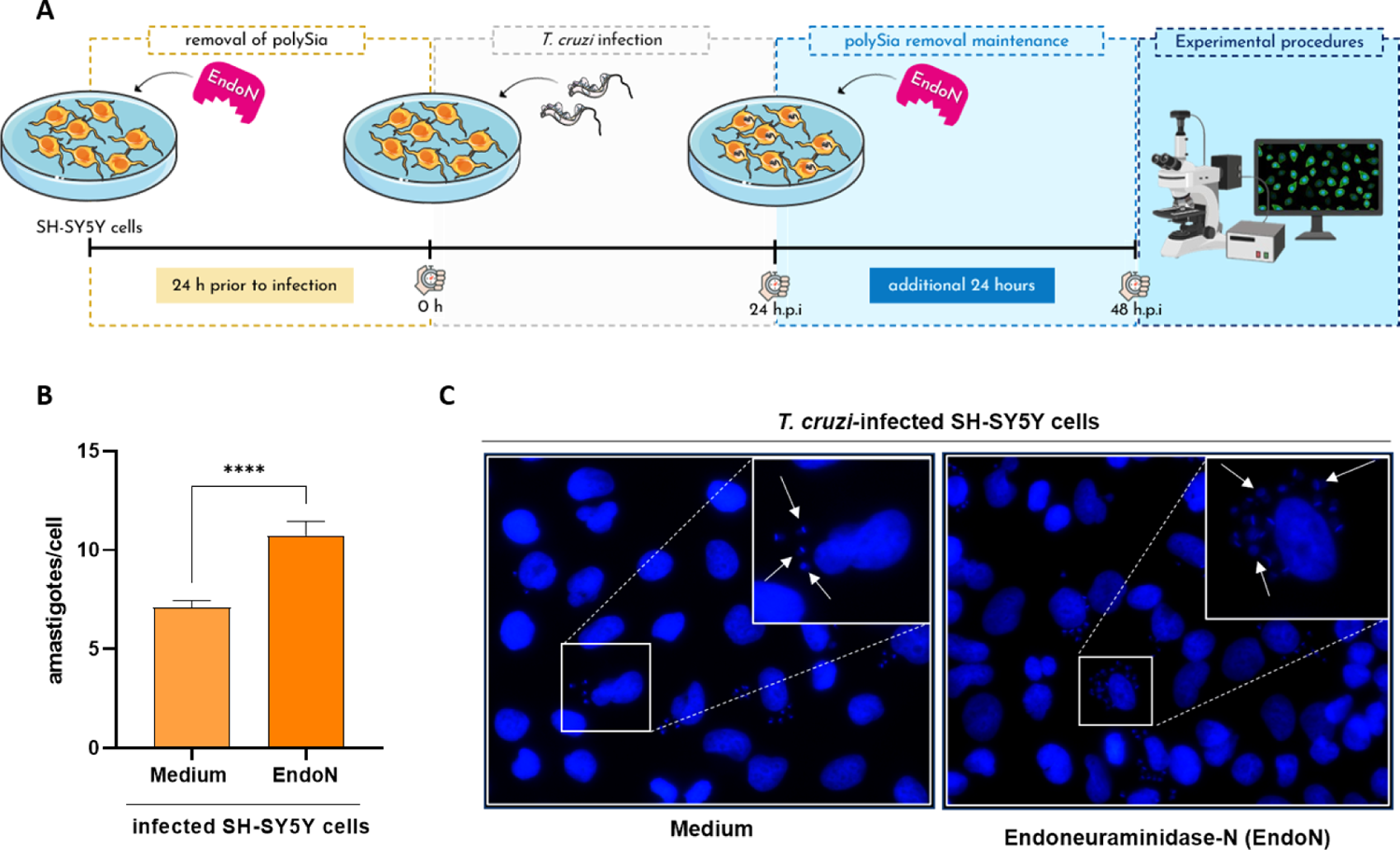
Removal of polySia in SH-SY5Y cells favors *T. cruzi* infection by increasing the number of internalized parasites. **(A)** Experimental workflow adopted to investigate the effect of enzymatic removal of polySia in *T. cruzi-*infected SH-SY5Y cells by EndoNeuraminidase (EndoN) [0.5 µg/ml]. SH-SY5Y cells were seeded in 24-well microplates (5 x 10^5^ or 5 x 10^4^ cells/well), and treated with EndoN [0.5 µg/ml] for 24h before infection, followed by addition of *T. cruzi* trypomastigotes (Y strain) at ratio of 1:5 (SH-SY5Y:trypomastigotes). After 24 h.p.i, *T. cruzi-*infected SH-SY5Y cells were submitted to another treatment with EndoN [0.5 µg/ml] for an additional 24h. Medium alone (Medium) were used as negative control; **(B)** Quantification of the number of intracellular parasites (amastigotes) per infected cell. Cell nucleus were stained with DAPI and following the experimental approach shown in **B**, amastigotes were counted in a total of 100 infected SH-SY5Y cells. Results are expressed as mean ± standard error of the mean (SEM) performed in triplicates; **(C)** Representative immunofluorescence images of *T. cruzi-*infected SH-SY5Y cells following the strategy adopted in the experimental workflow presented in **B**. Cell nucleus were stained with DAPI (scale bars = 10 µm). White arrows are representative of *T. cruzi* amastigotes inside in SH-SY5Y cells. Significant differences compared to the *T. cruzi-*infected SH-SY5Y treated with medium alone (Medium) are shown by (****) *p* < 0.0001; *ns*: not significant. Parts of the figure were drawn by using pictures from Servier Medical Art. Servier Medical Art by Servier is licensed under a Creative Commons Attribution 3.0 Unported License (https://creativecommons.org/licenses/by/3.0/), and BioRender.com under academic license.

### Chemical and genetic Inhibition of ST8Sia2 increases the number of parasites internalized in SH-SY5Y cells

The impact of *T. cruzi* infection on the modulation of host polysialylation was demonstrated here by different experimental approaches in Figures 1-4. Thereafter, we evaluated whether chemical and genetic inhibition of ST8Sia2 would have an impact on host cell infection by *T. cruzi*. For this, we used cytidine 5′-monophosphate (CMP), a molecule previously reported as a pharmacological inhibitor of ST8Sia2 [54], and the siRNA-directed approach. Initially, we demonstrated that chemical or genetic inhibition of ST8Sia2 in SH-SY5Y cells with CMP or siRNA ST8Sia2, respectively, does not affect cell viability **(Supplementary Figure S3A,B)**. Then, SH-SY5Y cells were treated with CMP and infected with *T. cruzi* according to the workflow shown in Figure 5A. The number of internalized parasites was determined after labeling them at 48 h post-infection. We demonstrated that chemical inhibition of ST8Sia2 with CMP in SH-SY5Y cells increased the number of intracellular parasites (Figure 5B,C). We also demonstrated that the number of intracellular parasites in untreated cells (culture medium only) or GMP-treated cells did not differ significantly (Figure 5D,E), demonstrating that the inhibition of ST8Sia2 favors infection by *T. cruzi*. Along the same purpose, we used a siRNA-directed approach to silence ST8Sia2 according to the workflow presented in Figure 5A.Transfection efficiency was evaluated by the relative expression of the ST8Sia2 transcript by qRT-PCR **(Supplementary Figure S3C**) according to the workflow shown in **(Supplementary Figure S3D**). Our results demonstrated that genetic silencing of ST8Sia2 in SH-SY5Y cells also increased the load of internalized parasites (Figure 5D**, E).** Taken together, these findings reinforce our hypothesis that ST8Sia2 inhibition favors *T. cruzi* infection.

**Figure 5.**
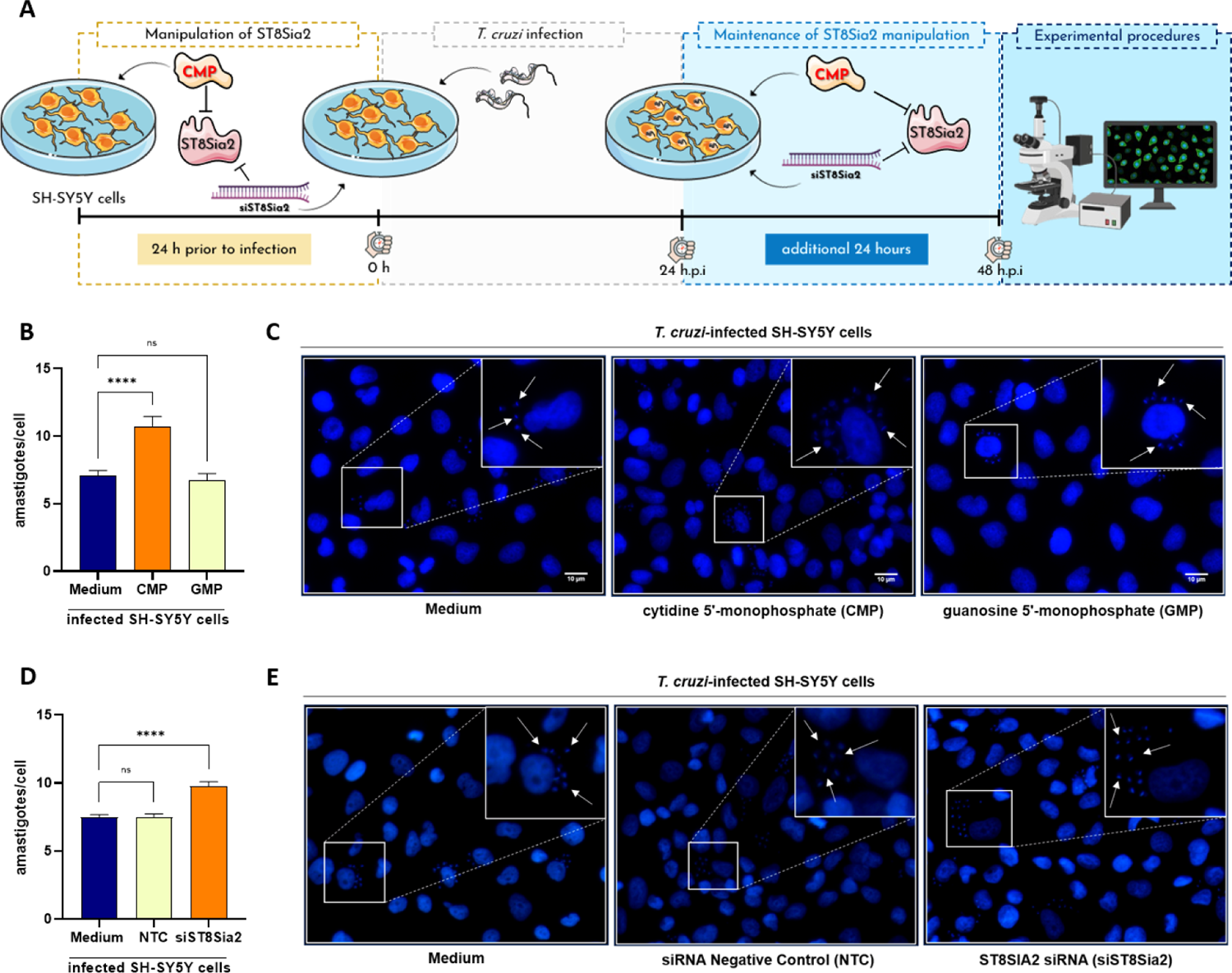
Chemical and genetic inhibition of ST8Sia2 sialyltransferase activity in SH-SY5Y cells favors *T. cruzi* proliferation. (A) Experimental workflow adopted to investigate the effect of chemical and genetic inhibition of ST8Sia2 in *T. cruzi-*infected SH-SY5Y cells using 0.5 mM of CMP. SH-SY5Y cells were seeded in 24-well microplates (5 x 10^5^ or 5 x 10^4^ cells/well), and previously treated with CMP [0.5 mM] or siRNA ST8Sia2 [100 nM] for 24h before infection, followed by infection with *T. cruzi* trypomastigotes (Y strain) at ratio of 1:5 (SH-SY5Y:trypomastigotes). After 24 h.p.i, *T. cruzi-*infected SH-SY5Y cells were treated with CMP [0.5 mM] or siRNA *ST8Sia2* [100 nM] for an additional 24h. Guanosine 5’-monophosphate (GMP), siRNA negative control (NTC), and/or medium alone (Medium) were used as negative controls for chemical or genetic inhibition of *ST8Sia2*; **(B,D)** Quantification of the number of intracellular parasites (amastigotes) per infected cell. Cell nucleus were stained with DAPI and following the experimental approach shown in **B**, amastigotes were counted in a total of 100 infected SH-SY5Y cells. Results are expressed as mean ± standard error of the mean (SEM) performed in triplicates; **(C,E)** Representative immunofluorescence images of *T. cruzi-*infected SH-SY5Y cells following the strategy adopted in the experimental workflow presented in **B**. Cell nucleus were stained with DAPI (scale bars = 10 µm). White arrows are representative of *T. cruzi* amastigotes inside in SH-SY5Y cells. Significant differences compared to the *T. cruzi-*infected SH-SY5Y treated with medium alone (Medium) are shown by (****) *p* < 0.0001; *ns*: not significant. Parts of the figure were drawn by using pictures from Servier Medical Art. Servier Medical Art by Servier is licensed under a Creative Commons Attribution 3.0 Unported License (https://creativecommons.org/licenses/by/3.0/), and BioRender.com under academic license.

## DISCUSSION

Different studies have extensively explored the manipulation of host cell glycosylation in infectious processes since many pathogens use this post-translational modification to ensure a successful infection [55,56]. In the case of *T. cruzi*, the role of Sia in the infection has been extensively explored, with a main focus on the transfer of this sugar from host α-2,3-sialylated glycans to the surface of the parasite catalized by trans-sialidase [37,40]. Also, knockout mice for CD43 [48], a sialoglycoprotein previously reported as a natural receptor for *T. cruzi* trans-sialidase [47], showed reduced cardiac parasite burning [48], reinforcing that manipulation of host glycosylation could directly or indirectly interfere in the *T. cruzi* infection. However, the impact of *T. cruzi* infection on host glycosylation machinery has not been explored to date.

Herein, we described the modulation of host cell N-glycosylation machinery in experimental *T. cruzi* infection, and, providing for the first time, evidences of downregulation of ST8Sia2, a host polysialyltransferase. Using an initial *in-silico* approach, different genes encoding enzymes involved in the protein glycosylation pathways, was suggested to be up (e.g. Neuraminidase 1 - Neu1, and α-1,3-mannosyltransferase - ALG3, or down (ST8 alpha-N-acetyl-neuraminide alpha-2,8-sialyltransferase 2 – ST8Sia2, and α-Glucosidase 2 - GANAB) regulated in the course of infection in human cardiomyocytes **(Supplementary Figure S1)** and in human fibroblasts (data not shown). Among these (glyco)genes, ST8Sia2 showed a sustained downregulation over different time points **(Supplementary Figure S1)**, which should lead at the end to a decrease in the polySia content of glycoproteins, as described in the literature [57].

Consistent with our initial observations, we found experimentally a significant reduction in *ST8Sia2* mRNA levels (Figure 1B) and polysialic content (Figure 1D) in human induced-pluripotent stem cells derived cardiomyocyte (hiPSC-CM) after 48 h of *T. cruzi* infection, although a less expressive decrease at the protein level of ST8Sia2 was observed at this time point (Figure 1C). Similar results, regarding decrease of ST8Sia2 expression (Figure 2B and C) and polySia (Figure 2B and D) were obtained with SH-SY5Y, a human neuroblastoma cell line, after 48 h of *T. cruzi* infection. A direct correlation between the expression of ST8Sia2 and the reduction in polySia levels has already been well demonstrated in previous studies that investigated the relevance of polysialylation in the nervous system [58,59].

Although polySia is added to N- and O-glycosylation structures, its presence is confined to a restricted set of glycoproteins, emphasizing NCAM1, which is the main and best-studied carrier of polySia in mammals [22], and in few other proteins, such as sodium voltage-gated channel alpha subunit 5 (SCN5A). Then, to better understand the impairment of host polysialylation in *T. cruzi* infection, we investigated whether NCAM1 and SCN5A would be differentially modulated in hiPSC-CM and SH-SY5Y during infection. In hiPSC-CM, NCAM1 levels did not differ between uninfected or infected cells (Figure 1E and G), in accordance with the observation that this cultured-cells do not express PSA-NCAM, the polysialylated form of NCAM1. On the other hand, the abundance of polySia in SCN5A, the primary cardiac isoform of the voltage-gated sodium channel (Na_v_ 1.5), was significantly reduced in hiPSC-CM-infected cells (Figure 1F and G). Interestingly, it was shown that differential sialylation in the primary cardiac isoform of Na_v_ 1.5 channel generates more depolarized potentials in atrial and ventricular cardiomyocytes, suggesting that cardiac contractility may be modulated by changes in sialic acids associated with the channels [51,60]. Furthermore, in ST8Sia2 knockout mice, polysialylation deficiency compromises action potential and Nav functions in atrial and ventricular cardiomyocytes [29,30]. Altogether, these results support our hypothesis that *T. cruzi* infection manipulates the polysialylation machinery of the host, compromising the expression of ST8Sia2, polySia, and SCN5A, which are targets of polysialylation, and thus may alter the complex biology of cardiomyocytes.

Similar to our findings with *T. cruzi-*infected hiPSC-CM, we detected reduced levels of ST8Sia2 and polySia in *T. cruzi-*infected human neuroblastoma (SH-SY5Y) cells, in addition to lower polysialylation of PSA-NCAM, expressed by this cell line (Figure 2B and E), or SCN5A (Figures 2 **and 3**). Metabolic differences between the cell lines employed may explain the lower levels in the polysialylated SCN5A and NCAM1, as well as in ST8Sia2 expression observed in SH-SY5Y in relation to hiPSC-CM cells (Figure 2 **and 3**). Nevertheless, both results reinforce our hypothesis that the polysialylation machinery is modulated during the course of *T. cruzi* infection. Although two different polysialyltransferases (ST8Sia2 and ST8Sia4) are involved in the synthesis of polySia chains, it was previously demonstrated that SH-SY5Y expresses high levels of *ST8Sia2* mRNA, but it is almost devoid of *ST8Sia4* mRNA [61] and expresses polySia bound to the two transmembrane proteoforms of NCAM, NCAM140, and NCAM180. Indeed, all expressed NCAM proteoforms are polysialylated [62], and at least, ST8Sia2 is related to polySia modulation of glycoproteins in cells infected by *T. cruzi*.

Although the role of NCAM is relatively little explored in the heart, its expression is known to be differentially regulated during the cardiac system development [63]. In the fetal heart, NCAM is expressed in the myocardium and ventricular conduction system, but in the adult heart, its expression is restricted to the ventricular conduction system [64]. Additionally, NCAM1 deletion compromises the expression of genes essential for the coordination of rhythmic contractions of the conduction system in a subpopulation of ventricular cardiomyocytes [29]. Considering that in chronic Chagas cardiomyopathy, the blockade of the ventricular conduction system by the high and persistent parasitism in the cardiac tissue leads to ventricular dysfunction, which can result in heart failure [65–67], it is tempting to speculate that somehow, the dysfunction in the ventricular conduction system frequently seen in chronic chagasic cardiomyopathy may result from manipulation of the polysialylation machinery that compromises PSA-NCAM and/or SCN5A expression.

Interestingly, both chemical or genetic inhibition of ST8Sia2 by CMP or siRNA *ST8Sia2*, respectively, increase the number of intracellular parasites in SH-SY5Y cells measured 48 h.p.i (Figure 4). This fact strongly suggests that in the course of the invasion of cells by *T. cruzi*, modulation of host polysialylation may be a strategy adopted by the parasite to ensure success in establishing the infection, either by affecting the invasion step (e.g. by decreasing the repulsive negative charges of surface molecules with time) or by affecting the intracellular cycle of the parasite. However, the mechanism, as yet, remains elusive. In this context, it is interesting to note that both, quantity and quality, of polySia synthetized by ST8SIA2 or ST8SIA4 are not so much different, but the resulted polySia-NCAMs exhibited different attractive and repulsive properties, which may affect biological functions [68]. To date, no study has explored the possible involvement of polysialylation in *T. cruzi* infection. However, previous studies reported that in Chagas disease, possible disorders of aberrant glycosylation are due to modifications in the sialylation of glycoproteins. As demonstrated before by Libby et al in 1986, *T. cruzi* trypomastigotes modify the surface of myocardial and vascular endothelial cells by desialylation [45].

In summary, the present study demonstrates the impairment of the host polysialylation machinery during infection by *T. cruzi*. We have shown that *T. cruzi* infection promotes the differential regulation of ST8Sia2 and polySia in host cells (figure 6). Using different experimental approaches, we also showed a reduction in NCAM and SCN5A, both targerts of ST8Sia2, which strongly suggests that ventricular conduction dysfunction frequently observed in chronic Chagas cardiomyopathy may be due to the manipulation of the polysialylation of these molecules. Finally, we demonstrate that pharmacological inhibition of ST8Sia2 in infected cells increased the number of intracellular parasites, thus providing a possible functional role for manipulating polysialylation during *T. cruzi* infection.

**Figure 6.**
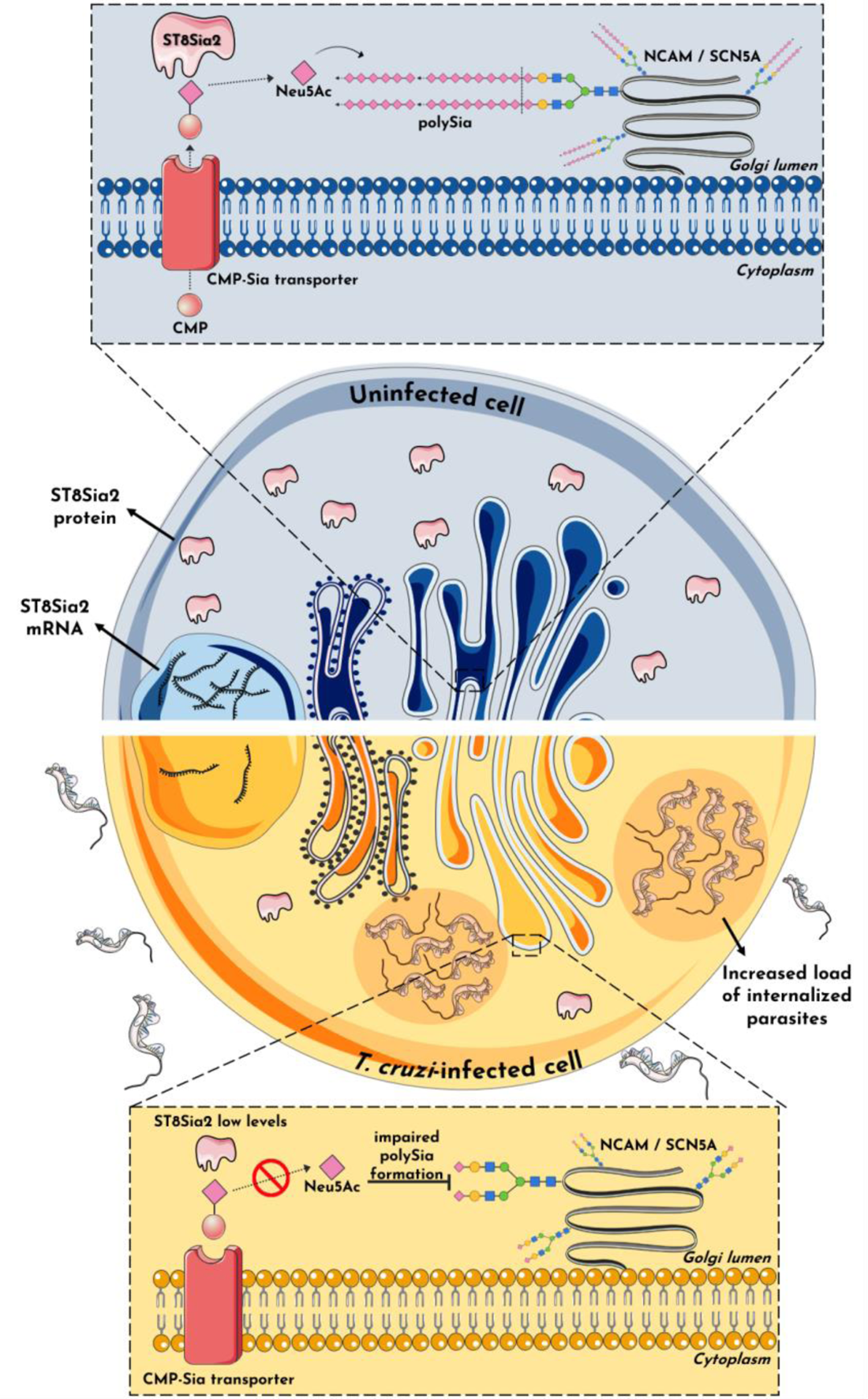
Hypothesized mechanism of host polysialylation modulation during *T. cruzi* Infection. The findings in this study suggest a significant impact on host polysialylation during *T. cruzi* infection. Under conditions of homeostasis, ST8Sia2 catalyzes the transfer of Neu5Ac monomers from cytidine monophosphate N-acetylneuraminic acid (CMP-Neu5Ac) to glycoproteins within the Golgi complex, forming a unique polymer of α-2,8-glycosidic linkages between Neu5Ac monomers known as polySia. This phenomenon, termed polysialylation, assumes pivotal roles across a spectrum of cellular functions. The addition of polySia into molecules such as NCAM1 and SCN5A significantly influences cardiac contractility and neural communication, exemplifying its multifaceted significance. In this study, we present, for the first time, evidence suggesting that host polysialylation exerts a substantial impact on *T. cruzi* infection dynamics. Our investigation has unveiled that *T. cruzi* infection profoundly diminishes the expression/abundance of the ST8Sia2 enzyme, compromising polySia formation, and its subsequent addition to polysialylated target molecules, including SCN5A and NCAM1. Consequently, the attenuation of ST8Sia2 and polySia levels may compromise the host’s ability to resist *T. cruzi* infection, as polysialylation modulation is accompanied by an increase in the number of internalized parasites. These findings strongly suggest that ST8Sia2 and, by extension, polySia, constitute a protective shield against *T. cruzi* infection. Their diminishment notably facilitates the establishment of infection. It is plausible to posit that the modulation of host polysialylation by the parasite serves as a pivotal determinant in the pathogenesis of Chagas disease. Parts of the figure were drawn by using pictures from Servier Medical Art. Servier Medical Art by Servier is licensed under a Creative Commons Attribution 3.0 Unported License (https://creativecommons.org/licenses/by/3.0/).

## MATERIALS AND METHODS

### Public RNA-seq data reanalysis

The fastq files were downloaded from the https://sra-explorer.info/ platform with the BioProject accession number PRJNA532430 [49]. The ‘FastQC’ module was used to report the quality reads, followed by the trimmed of adapters using TrimGalore! set to the single-end library. The TrimGalore! output sequences were aligned to the human reference genome hg38 using the HISAT2 platform [69]. A count table was generated using the FeatureCounts [70] algorithm. The differently regulated genes were analyzed by the limma, Glimma, edgeR, and Homo.sapiens packages applying a cut-off of |log2FC|>1 and a p-adjusted value <0.05 (Benjamini-Hochberg). Statistical data of the glycogenes of interest were filtered and the chord_dat() function, available in the GOplot package, was used to plot the chord plot. Volcano plot and heatmap graphics were made using Perseus software.

### Human cell culture

Human induced-pluripotent stem cells (hiPSC) were differentiated into Human induced-pluripotent stem cells derived cardiomyocyte (hiPSC-CM) on Geltrex (Thermo Fischer, Waltham, MA, USA) coated plates in Cardiomyocytes-Plating Medium (Pluricell Biotech), using an established differentiation protocol [71]. All experiments were carried out between day 30-35 of differentiation. The human neuroblastoma SH-SY5Y cells (ATCC-CRL-2266) were cultured in Dulbecco’s Modified Eagle Medium/Nutrient Mixture F-12 (DMEM/F12) (Gibco®, Life Technologies, Carlsbad, CA, USA), supplemented with 10% heat-inactivated fetal bovine serum, and 1% penicillin/streptomycin. All cells line were maintained in a humidified 5% CO_2_ atmosphere at 37 °C.

### *T. cruzi* culture and infection assay

*T. cruzi* trypomastigote (Y strain) was routinely maintained by infection in monkey kidney epithelial cells (LLC-MK2) in Roswell Park Memorial Institute 1640 Medium (RPMI) supplemented with 2% fetal bovine serum in a humidified 5% CO_2_ atmosphere at 37 °C, according to previously described experimental procedures [72]. Five days after infection, the trypomastigotes released in the supernatants were purified using a DEAE-cellulose chromatography column. Therefore, the cells were washed in phosphate-buffered saline (PBS) by centrifugation at 10.000 × g for 12 min, and resuspended in adequate culture medium for each human cell line used. The total number of parasites/ml was counted in a Neubauer chamber. hiPSC-CM and/or SH-SY5Y cells were seeded at a density of 5×10^5^ cells/well in 24-well or 6-well microplates and infected with purified trypomastigotes at the rate of 5:1 (trypomastigotes:hiPSC-CM) in a volume of 500 μL per well [49]. After 3 h of incubation, the media was replaced with fresh media to remove the non-internalized parasites. The impact of *T. cruzi* infection on host polysialylation was evaluated at 48 h post-infection (h.p.i). In all experiments, an uninfected group was subjected to the same experimental conditions, except for incubation with *T. cruzi*.

### Immunofluorescence assay

hiPSC-CM or *T. cruzi-*infected SH-SY5Y cells distributed in round glass coverslips (13 mm) contained in 24-well microplates with flat bottom were fixed with PBS/4% paraformaldehyde for 15 min at room temperature (RT) and permeabilized with 0.1% Triton X-100 (Sigma, Chemical Company, St. Louis, MO, USA) for 15 min at RT. For evaluation the efficiency of differentiation on hiPSC-CM, coverslips were blocked with PBS/2% Bovine Serum Albumin (BSA) solution, and incubated with primary antibodies: mouse anti-cardiac troponin T (HyTest clone 4T19/2) and rabbit anti-α-actinin (Millipore) for 30 min at RT. After washing four times with PBS, the cells were incubated for 1 h at RT with the secondary antibodies: goat anti-rabbit immunoglobulin G (IgG) Alexa Fluor 488 and/or goat anti-mouse IgG Alexa Fluor 594 (Thermo Fisher Scientific, Waltham, Massachusetts, USA). The coverslips were washed four times and prepared for visualization by adding one drop of mounting medium (Slow Fade Gold Antifade; Thermo Fisher Scientific) with 4’,6-diamidino-2-phenylindole (DAPI). For evaluation of the ST8Sia2 and polySia levels in *T. cruzi-*infected SH-SY5Y cells, the cells seeded on coverslips, fixed and permeabilized were blocked with PBS/2% Bovine Serum Albumin (BSA) solution, and incubated with primary antibodies: ST8Sia2 monoclonal antibody (Catalog WH0008128M1, Sigma-Aldrich, St. Louis, MO, USA) and polySia-specific monoclonal antibody (mouse IgG2a, clone 735) for 30 min at RT. After washing four times with PBS, the cells were incubated for 1 h at RT with the secondary antibodies: Alexa Fluor^TM^ 488 goat anti-mouse IgG (Thermo Fisher Scientific, Waltham, Massachusetts, USA) or Alexa Fluor^TM^ 594 goat anti-mouse IgG (Thermo Fisher Scientific, Waltham, Massachusetts, USA). The coverslips were washed four times and prepared for visualization by adding one drop of mounting medium (Slow Fade Gold Antifade; Thermo Fisher Scientific) with 4’,6-diamidino-2-phenylindole (DAPI). To determine the number of parasites internalized in SH-SY5Y cells, the cells seeded on coverslips, fixed and permeabilized as above, were prepared for visualization by adding a drop of mounting medium (Slow Fade Gold Antifade; Thermo Fisher Scientific) containing 4’,6-diamidino-2-phenylindole (DAPI), used for imaging, which was performed using a Conventional Fluorescence Microscope (Zeiss). polySia-specific monoclonal antibody (mouse IgG2a, clone 735) were kindly provided by Dr. Martina Mühlenhoff (Institute of Clinical Biochemistry, Hannover Medical School, Germany).

### Quantification of immunofluorescence

Fluorescence microscopy images were analyzed by ImageJ software, version 1.49v (Rasband, W.S., ImageJ, U. S. National Institutes of Health, Bethesda, Maryland, USA, http://imagej.nih.gov/ij/) to evaluate ST8Sia2 and polySia-stained host cells. Using the calculation for corrected total cell fluorescence (CTCF) = integrated density–(area of selected cell × mean fluorescence of background readings), as described by Gachet-Castro et al. [73]. The fluorescence intensity of each cell was calculated. Subsequently, graphs were plotted with the fluorescence average values using GraphPad Prism 9.0 program (GraphPad Software, Inc., San Diego, CA).

### Total Cell Lysates

hiPSC-CM and SH-SY5Y cells were infected with *T. cruzi* trypomastigotes at the rate of 5:1 (trypomastigotes:hiPSC-CM) for 48 h. After 48 h.p.i, the cells were washed four times with PBS to remove extracellular parasites, and lysed with radioimmunoprecipitation assay (RIPA) buffer [20 mM Tris pH 7.2, 150 mM NaCl, 1% Triton X-100, 1% sodium deoxycholate, 0.1% sodium dodecyl sulphate (SDS)] containing a cocktail of protease (cOmplete, Sigma-Aldrich, St. Louis, MO, USA) and phosphatase (PhosStop, Sigma-Aldrich) inhibitors. The cell lysates were kept on ice for 10 min, centrifuged at 14.000 x g for 10 min at 4 °C to pellet cell debris. Supernatants were collected, and the total protein was measured using the Kit Pierce^TM^ BCA Protein Assay (Thermo Scientific) according to the manufacturer’s instructions. In all experiments, an uninfected group was subjected to the same experimental conditions. All Samples were resuspended in sample buffer for analysis by SDS-PAGE. The protein extract from human rhabdomyosarcoma cell line TE671 in sample buffer [2% sodium dodecyl sulfate, 10% glycerol, 0.002% bromophenol blue, 0.0625 M Tris-HCl pH 6.8, 5% 2-mercaptoethanol] were gifts from Dr. Martina Mühlenhoff (Institute of Clinical Biochemistry, Hannover Medical School, Germany).

### Immunoblotting assay

Proteins extracted from total cell lysate (15μg of proteins) were resolved by SDS-PAGE and transferred to PVDF membranes, which were directly incubated with blocking buffer (5% BSA in Tris-buffered saline (TBS) at 0.05% Tween-20 (TBS/T) for 1 h. Subsequently, membranes were washed three times with TBS/T solution and incubated *overnight* with the primary antibodies: anti-ST8Sia2, anti-polySia, anti-NCAM1, anti-PSA-NCAM, anti-SCN5A, and Anti-ACTB, and washed three times with TBS/T. Then, the membrane was incubated with the respective secondary antibodies for 1 h at RT, followed by the incubation with a secondary antibody for 1h at room temperature. The immunoreactive bands were detected with the ChemiDoc XRS Imaging System equipment and protein quantification was performed using the ImageJ software. polySia-specific monoclonal antibody (mouse IgG2a, clone 735) and NCAM1-specific monoclonal antibody (mouse IgG1, clone 123C3) were kindly provided by Dr. Martina Mühlenhoff (Institute of Clinical Biochemistry, Hannover Medical School, Germany). All antibodies used in this study were tested for crossreactivity against *T. cruzi* trypomastigote cell lysates.

### Drugs treatment and chemical ST8Sia2 inhibition

SH-SY5Y cells were pretreated with 0.5 mM cytidine 5’-monophosphate (CMP, Sigma-Aldrich) or culture medium (negative control) 24 h before infection and 24 h post-infection with *T. cruzi* to evaluate the impact of pharmacological inhibition of ST8Sia2 activity during infection and replication of the parasite. After incubation with CMP for 24 h, the CMP-containing medium was replaced with fresh medium, and the cells were infected with *T. cruzi* trypomastigotes in a 5:1 ratio (trypomastigotes:hiPSC-CM). 24 h post-infection, the treatment with 0.5 mM CMP was repeated for another 24 h. Guanosine 5′-monophosphate (GMP), with no effect on ST8Sia2 inhibition, was used as a control. In all experiments, a group of infected cells (negative control of ST8SIa2 inhibition) was subjected to the same experimental conditions, except for incubation with pharmacological inhibitor.

### siRNA-Directed Inhibition of ST8Sia2

Predesigned siRNAs for the *ST8Sia2* transcripts (*siST8Sia2*) (hs.Ri.ST8Sia2.13.2; hs.Ri.ST8Sia2.13.3) were purchased from Integrated DNA Technologies (IDT, Coralville, IA). Transfection was performed using Lipofectamine RNAiMAX (Thermo Fisher Scientific, Waltham, MA, USA). Lipofectamine RNAiMAX:siST8Sia2 Mix (0.6 µL Lipofectamine:100 nM siRNAs) was formed in Opti-MEM (Thermo Fisher Scientific, Waltham, MA, USA) at room temperature for 20 min and added to SH-SY5Y cells in antibiotic-free medium for overnight transfection at 37 °C and 5% CO_2_. After incubation, fresh Dulbecco’s Modified Eagle Medium/Nutrient Mixture F-12 (DMEM/F12) (Gibco®, Life Technologies, Carlsbad, CA, USA), supplemented with 2% heat-inactivated fetal bovine serum was added, and the cells were infected with *T. cruzi* trypomastigotes in a 5:1 ratio (trypomastigotes:hiPSC-CM). 24 h post-infection, *T. cruzi-*infected SH-SY5Y cells were transfected again with Lipofectamine RNAiMAX:siST8Sia2 Mix (0.6 µL Lipofectamine:100 nM siRNAs) in Opti-MEM (Thermo Fisher Scientific, Waltham, MA, USA) antibiotic-free medium for an additional 24 h, at 37 °C and 5% CO_2_. siRNA negative control (NTC) and medium alone (Medium) were used as negative controls. Transfection efficiency was evaluated by the relative expression of the ST8Sia2 using Quantitative real-time reverse transcription PCR (qRT-PCR).

### Quantitative real-time reverse transcription PCR (qRT-PCR)

hiPSC-CM were infected with *T. cruzi* trypomastigotes (Y strain) at the rate of 10:1 or 5:1 (trypomastigotes:host cell). Total RNA from these samples was extracted using TRIzol according to the manufacturer’s instructions, and the RNA was converted into cDNA using the High-Capacity cDNA Reverse Transcription kit (Applied Biosystems™). qRT-PCR was performed in a final volume of 10 μL using Maxima™ SYBR Green qPCR Master Mix (Thermo Scientific) and the quantification was done by QuantStudio™ 3 Real-Time PCR System (Applied Biosystems™). The cycling conditions were as follows: initial denaturation at 95 °C for 10 min, followed by 40 cycles of denaturation at 95 °C for 15 s and annealing/extension at 60 °C for 60 s. The relative expression of the transcript was quantified using the ΔΔCt method [74] and normalized to β-actin expression. The specificity of amplification was determined using melting curve analysis. The following PCR primers were used: β-actin (F-CATGTACGTTGCTATCCAGGC, R-CTCCTTAATGTCACGCACGAT), and ST8Sia2 (F-CGTTCGCGGATACTGGCT, R-GCCAGAAGCCGTAGAGGTAG).

### MTT assay

SH-SY5Y cells were seeded in 96-well microplates (1 × 105 cells/well), and treated with cytidine 5’-monophosphate (CMP; 62.5 – 500 µM) or siRNA ST8Sia2 (50 – 150 nM) for 24 and/or 48 h at 37 °C. Guanosine 5’-monophosphate (GMP), siRNA negative control (NTC), and/or medium alone were used as negative controls for cell death. The cell viability of the cells was determined after the reduction of MTT (Sigma-Aldrich) to produce formazan crystals [75]. The procedure was performed as described by Barboza *et al.* [3]. Mitochondrial activity was expressed as a percentage after comparison with the absorbance of unstimulated SH-SY5Y cells.

### Quantification of polysialic acid

For quantification of polysialic acid, SH-SY5Y cells infected with *T. cruzi*, as well uninfected cells, were incubated with iEndo-N (inactive Endo-N-acetylneuraminidase) for 1h at 37°C. After the incubation time, 30 µg of protein from each sample were immobilized on PVDF membrane and submitted to direct hydrolysis of the sialic acid with 2M acetic acid for two hours at 80° C. Hydrolyzed Sia were derivatized with 1,2-Diamino-4,5-methylenedioxybenzene (DMB; P/N D4784) obtained from Sigma-Aldrich and separated in reversed-phase chromatography with fluorescence detection using a Vanquish UPLC (Thermo Fisher). iEndo-N was a gift from Dr. Martina Mühlenhoff (Institute of Clinical Biochemistry, Hannover Medical School, Germany).

### Statistical analysis

Initially, the results were tested for normal distribution and homogeneous variance. Once the normal distribution and homogeneous variance was confirmed, the One-way ANOVA test was applied. In the absence of normal distribution and heterogeneous variance, the Kruskal-Wallis test was applied. The statistical evaluation of the differences between the means of the groups were determined by the one-way analysis of variance, followed by the Bonferroni multiple comparison test and, in the non-parametric test, the comparisons between the groups were carried out by the Dunn’s multiple comparison test. For all statistical analyzes, the Prism 9.0 program (GraphPad Software, La Jolla, CA, USA) was used.

## ACKNOWLEDGMENTS

We thank Celia Ludio Braga at the Department of Biochemistry, Institute of Chemistry, University of São Paulo, Brazil for helping with the parasite cell culture. We thank Dr. Claudia Blanes Angeli at the GlycoProteomics Laboratory, Department of Parasitology, Institute of Biomedical Sciences, University of São Paulo, Brazil, for technical support.

## SUPPORTING INFORMATION

**Supplementary Figure S1.**
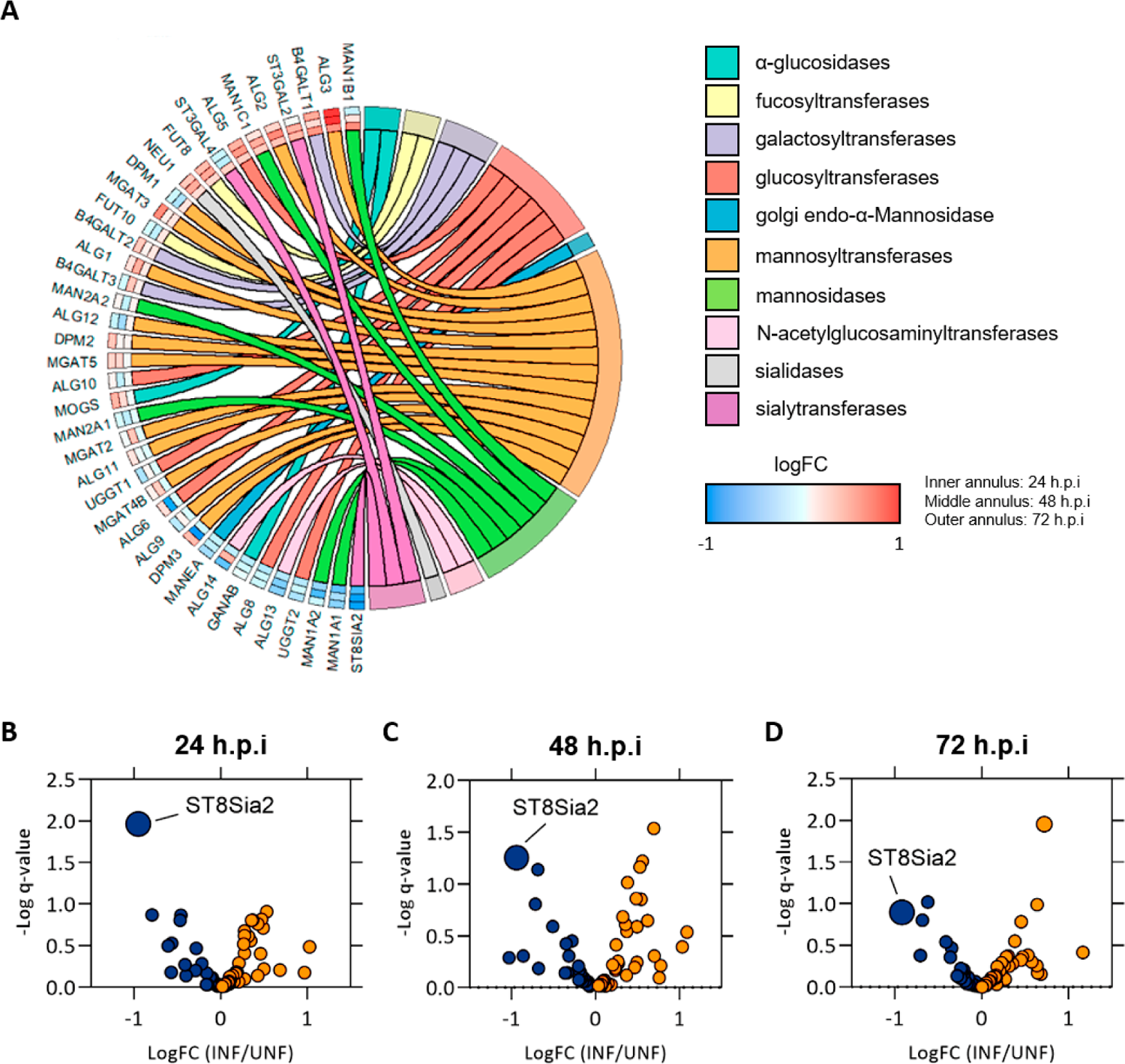
*T. cruzi* infection modulates genes involved in host N-glycosylation machinery. Bioinformatics analysis of Human induced pluripotent stem cell-derived cardiomyocytes (hiPSC-CM) infected with *Trypanosoma cruzi* (Y strain). **(A)** Chordplot indicating the modulation of N-glycosylation machinery transcripts at 24 h (inner), 48 h (middle), and 72 h (outer) post-infection (h.p.i) with *T. cruzi* (infected vs non-infected; INFvsUNF). The R GOplot package was used to build chord plot. Red and blue colors indicate upregulated and downregulated transcripts, respectively; **(B-D)** Volcano plot indicating the modulation of N-glycosylation pathway transcripts focusing on the modulation of the ST8Sia2 on *T. cruzi-*infected hiPSC-CM after 24 h **(B)**, 48 h **(C)**, and 72h **(D)** post-infection. The logFC (INFvsUNF) indicates transcripts upregulated in orange and downregulated in dark blue.

**Supplementary Figure S2.**
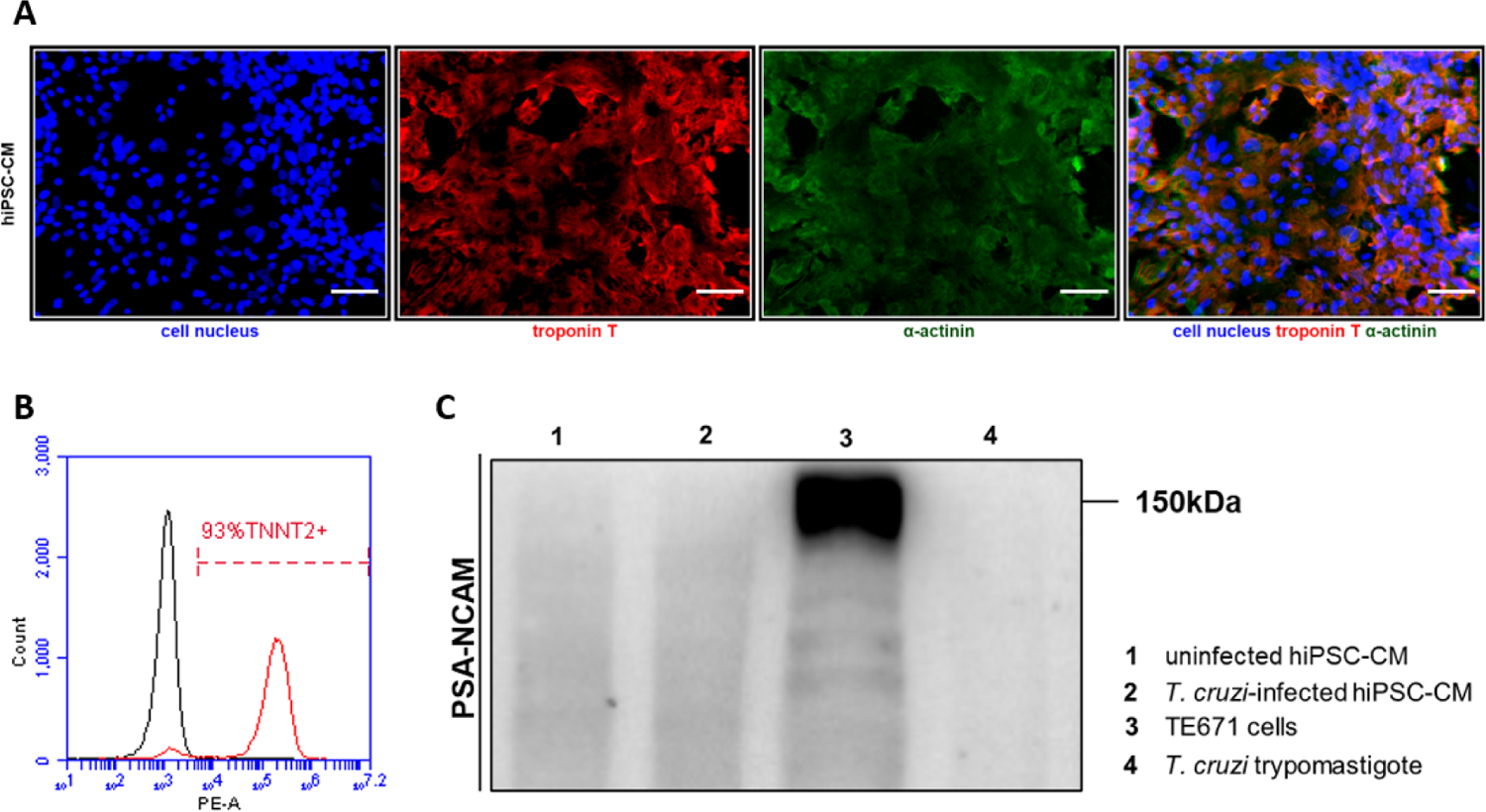
Human pluripotent stem cells (hiPSC) are efficiently differentiated into cardiomyocytes (hiPSC-CM), and do not express PSA-NCAM. (A) Representative immunofluorescence images of hiPSC-CM confirming the presence of α-actinin and troponin T (sarcomeric proteins). Cell nucleus were stained with DAPI (scale bars = 10 µm). (B) Histogram of Troponin T (TNNT2+) staining the differentiation of hiPSC into hiPSC-CM. (C) Representative western blot images of PSA-NCAM levels in uninfected hiPSC-CM (lane 1), *T. cruzi-*infected hiPSC-CM (lane 2), TE671 cells (lane 3), and *T. cruzi* trypomastigote (lane 4). hiPSC-CM cells were seeded in 6-well microplates (1 x 10^6^ cells/well) and infected with *T. cruzi* trypomastigotes (Y strain) in a ratio of 1:5 (SH-SY5Y:trypomastigotes) for 48 h at 37 ° C. Protein extracts from TE671 cell lysates (in Laemmli buffer) and *T. cruzi* trypomastigote (in RIPA buffer) were used as a positive and negative control for PSA-NCAM, respectively. 15 µg of protein extracts were used to analyze the PSA-NCAM levels by Western blotting using a specific anti-PSA-NCAM antibody.

**Supplementary Figure S3.**
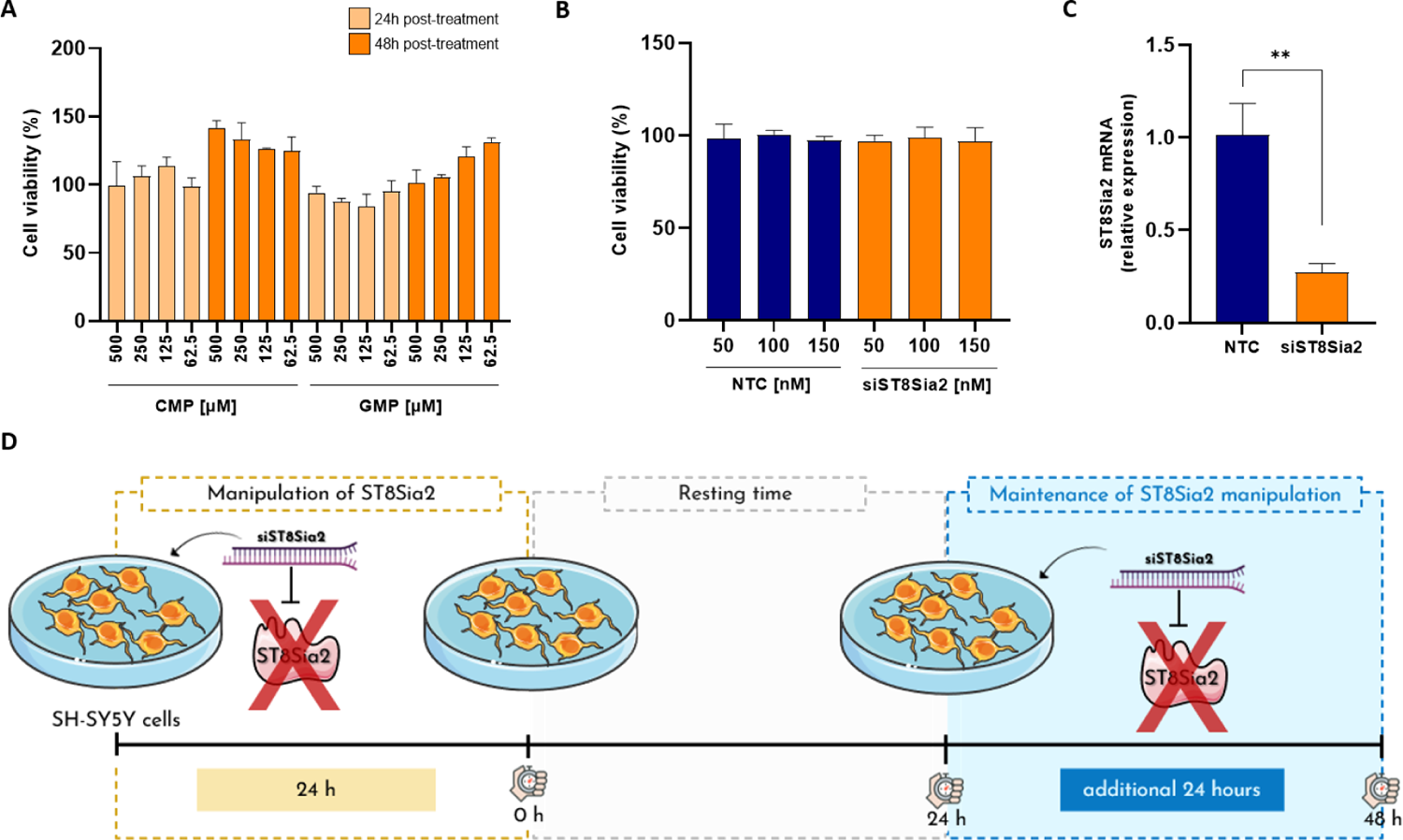
Chemical and genetic inhibition of ST8Sia2 in SH-SY5Y cells does not affect cell viability. **(A,B)** SH-SY5Y cells were seeded in 96-well microplates (1 × 10^5^ cells/well), and treated with cytidine 5’-monophosphate (CMP; 62.5 – 500 µM) or siRNA *ST8Sia2* (50 – 150 nM) for 24 and/or 48 h. Guanosine 5’-monophosphate (GMP), siRNA negative control (NTC), and/or medium alone were used as negative controls for chemical or genetic inhibition of ST8Sia2. Following 24 and 48 h incubation with CMP **(A)** or 24 h incubation with siRNA *ST8Sia2* **(B)**, 3-(4,5-dimethyl-2-thiazolyl)-2,5-diphenyl-2H-tetrazolium bromide (MTT; 50 μg/mL) was added to the cells, and mitochondrial activity was estimated by MTT reduction and expressed as a percentage calculated from the ratio between the absorbance of stimulated and non-stimulated SH-SY5Y cells; **(C)** Relative expression of *ST8Sia2* mRNA measured by qRT-PCR in SH-SY5Y cells treated with siRNA *ST8Sia2* (*siST8Sia2*), following the experimental workflow presented in **D**. siRNA negative control (NTC) were used as negative control for genetic silencing of ST8Sia2. The Ct values of the target transcripts were normalized to the relative expression of *GAPDH* as endogenous control, and the relative expression of ST8Sia2 transcripts was quantified by the 2^-ΔΔ^ Ct method. Each bar represents the mean of three independent experiments performed in triplicate; **(D)**. Experimental workflow adopted to investigate the effect of genetic inhibition of ST8Sia2 in SH-SY5Y cells using 100 nM of siRNA *ST8Sia2*. SH-SY5Y cells were seeded in 24-well microplates (5 x 10^4^ cells/well), and treated with siRNA *ST8Sia2* [100 nM] for 24h, followed by resting time of 24 h additional. After this resting time, siST8Sia2 SH-SY5Y cells were restimulated with siRNA *ST8Sia2* [100 nM] for an additional 24h. siRNA negative control (NTC) were used as negative control. Significant differences compared to the *siST8Sia2* SH-SY5Y cells with siNTC SH-SY5Y cells are shown by (**) *p* < 0.05; *ns*: not significant. Parts of the figure were drawn by using pictures from Servier Medical Art. Servier Medical Art by Servier is licensed under a Creative Commons Attribution 3.0 Unported License (https://creativecommons.org/licenses/by/3.0/).

